# Exploring *Mycobacterium tuberculosis* rRNA Transcriptomic Signatures as Response to Anti-TB Treatment in Whole Blood RNA-seq Data

**DOI:** 10.1101/2025.06.16.659901

**Authors:** Patricio Lopez-Exposito, Juan Calvet Seral, Santiago Ferrer, Alfonso Mendoza, Pedro M. Gordaliza, Juan Jose Vaquero

## Abstract

Current clinical tools for monitoring tuberculosis (TB) treatment response rely mostly on sputum culture and chest X-ray, which are unreliable in patients with low bacterial loads and lack the desirable promptness. There is a need for biomarkers able to provide earlier and more accurate insights into pathogen viability and response to therapy. We analyzed publicly available whole-blood RNA-seq data from 79 TB patients sampled at diagnosis, and weeks 1, 4, and 24 of standard anti-TB treatment. After aligning human reads and filtering, remaining reads were mapped to the *Mycobacterium tuberculosis* H37Rv genome to quantify rRNA subunit transcripts (16S, 23S, 5S, ITS1, ITS2). Microbiome profiles were assessed using Kraken2/Bracken, with alpha/beta diversity analyses and differential abundance (ANCOM-BC2). 16S and 23S rRNA transcripts were consistently detected across all treatment times, with 23S reads dominating in diagnosis and early stages of treatment shifting toward a significant predominance of 16S reads at week 24. 5S and ITS1 were inconsistently detected, whereas ITS2 was undetectable. Alpha diversity (Shannon index) increased during treatment (significant at weeks 1 and 4), while beta diversity showed significant shifts over time despite no significant differences in total *M. tuberculosis* abundance. Our findings suggest that it may be feasible to detect *M. tuberculosis* rRNA signatures in blood RNA-seq and suggest dynamic transcriptomic changes during treatment. The 16S/23S ratio and minor rRNA units may serve as complementary biomarkers for treatment monitoring and other transcriptomic-based biomarkers. Future work should validate these findings in larger cohorts using optimized RNA-seq protocols focusing on the pathogen.

## Introduction

Effective monitoring of treatment response in tuberculosis (TB) patients remains a critical challenge in clinical practice. Current approaches to stratify patients—such as clinical assessment, radiological imaging, and sputum culture conversion—often require several weeks to months to detect treatment failure. These methods are particularly limited in patients with difficulties producing sputum, and in those with low bacterial loads or extrapulmonary tubercular infection, delaying crucial treatment adjustments (Goletti et al., 2018). While molecular diagnostic tools like Xpert MTB/RIF have improved the speed and accuracy of initial drug resistance detection (WHO, 2021; Nathavitharana et al., 2017), they only provide a static snapshot of *Mycobacterium tuberculosis* (Mtb) at baseline, offering no dynamic insight into pathogen viability or real-time treatment response. The development of complementary biomakers capable of tracking bacterial activity over the course of therapy would be desirable to improve treatment efficacy and reduce patients suffering.

To address these limitations, recent works have explored pathogen-based transcriptomic signatures, including the Molecular Bacterial Load Assay (MBLA) as a surrogate for viable bacterial load (Honeyborne et al., 2014) and the ratio of precursor to mature *M. tuberculosis* rRNA (RS ratio) as an index of drug action that distinguishes between bactericidal and sterilizing effects (Walter et al., 2021). As can be derived from above, ribosomal RNA (rRNA) transcripts, in particular, represent promising biomarkers due to their high abundance, stability, and sensitive response to metabolic and translational activity. Drug preesure alters rRNA biosynthesis and physiology (Laureti et al. 2013, Briffotaux et al. 2019), what may translate into changes in the relative expression of rRNA subunits—such as 16S, 23S, and 5S—that could serve as early indicators of treatment impact.

Building on this hypothesis, we investigate whether rRNA-derived signatures of *M. tuberculosis* can be extracted from host-filtered whole-blood RNA-seq data and quantified longitudinally throughout standard TB treatment. In particular, we assess the detectability and dynamics of major rRNA transcripts at different time points and explore their potential as molecular markers of treatment response. By re-purposing public transcriptomic data from TB patients undergoing therapy, our study aims to evaluate the feasibility of using rRNA expression profiles as a novel class of non-invasive, pathogen-derived treatment monitoring biomarkers that can be complementary to current established tools or to those in development.

## Materials and Methods

### Dataset

RNA seq data (SRA study <u>SRP092402</u>) were obtained from a longitudinal study (Odia et al, 2012), in which whole blood samples were extracted from pulmonary TB patients at various timepoints. All patients were diagnosed with active tuberculosis (ACTB) through microbiological sputum culture examination. Patients were between 16 and 70 years old, had no other lung disease, were non-diabetic, and were HIV negative at diagnosis. All patients were treated according to WHO guidelines, that is, combination therapy for 6 months of combination therapy comprising two phases, the first consisting of two months of daily administration of rifampicin, isoniazid, pyrazinamide and ethambutol, followed by a second 4 month phase with rifampicin and isoniazid daily. We selected those elements from the whole dataset which presented all time points, namely diagnosis, week 1, week 4 and week 24 (N=79). Blood samples were extreacted employing Paxgene RNA extraction kits and latter to GlobinClear Globin transcript reduction followed by Illumina mRNA-Seq.

### Analysis pipeline

The pipeline for analysis is schematized in Figure 1. We used STAR 2.5.2b to align the data to the human genome GRCh38 and filtered out non-human reads. We then aligned non-human reads to the *Mycobacterium tuberculosis* H37Rv reference genome ((Cole & Barrell, 1998)), employing BWA-MEM2 2.2.1. We used Samtools v1.21 to count the mappings to the main regions of the Mtb ribosomal RNA operon, namely 16S, 23S and 5S and the corresponding internal transcribed spacer regions ITS1 and ITS2. In parallel, the microbiome analysis of the samples was performed with Kraken2 v2.0.8 by means of the PlusPF-16 pre-built database. Bracken v2.8 was employed to refine the output of Kraken2. Alpha and beta diversity analysis were performed employing scikit-bio v0.6.3 (Rideout et al., 2023). We applied PERMANOVA to the Bray–Curtis distance matrix stratified by subject using the adonis2 function from vegan v2.7-1 R package. Differential abundance of the subsequent outcome were analysed with R package ANCOM-BC2 v2.2.2. Between-groups comparison used Wilcoxon signed-rank with Benjamini-Hochberg correction for multiple testing.

**Figure 1.**
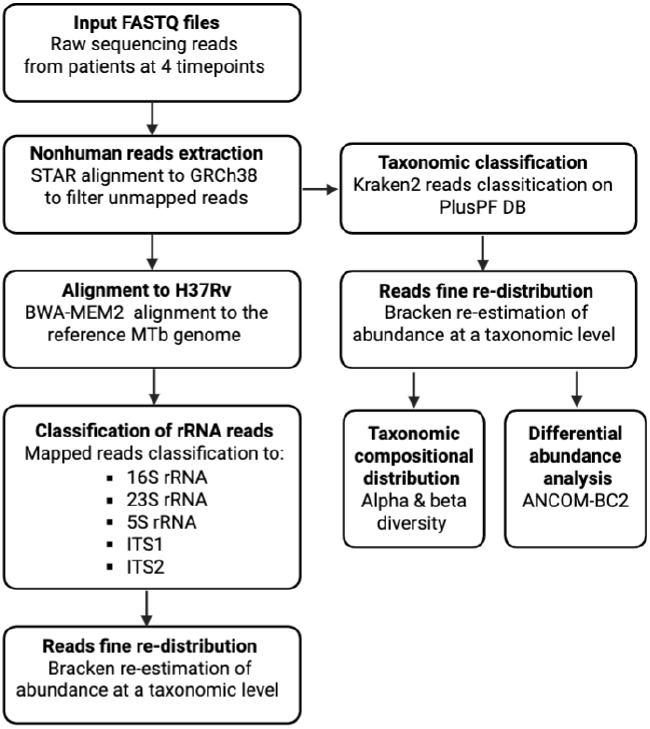
pipeline for processing whole-blood RNA-seq data and quantifying *M. tuberculosis* rRNA transcripts during anti-TB treatment.

## Results

### Distribution of rRNA transcripts in the course of the treatment

The average number of reads associated with the rRNA units per treatment time over all patients is shown in Figure 2. Transcripts corresponding to 16S and 23S units were found in all samples, whereas 5S and ITS1 transcripts were detected in some of them. ITS2 was not detected in any sample. Figure 3.a depicts the proportion of transcripts corresponding to units 16S and 23S relative to the all transcripts mapped to the Mtb rRNA operon.

**Figure 2.**
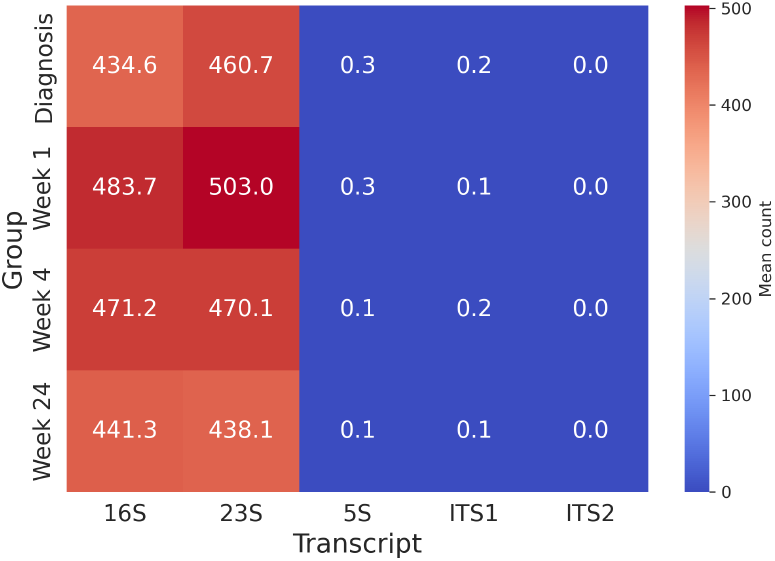
Total counts per treatment time and rRNA sub-unit.

**Figure 3.**
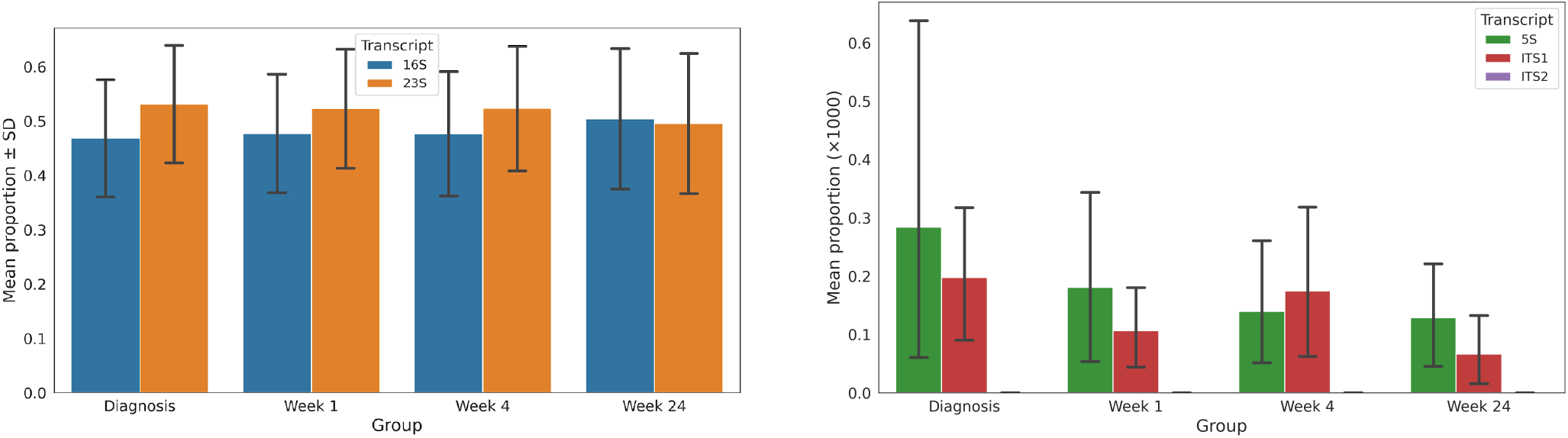
a) Proportion of 16S and 23S transcripts per treatment time and b) Scaled proportions of 5S and ITS1 subunitsfor each timepoint

In Figure 3.a the proportions of major rRNA 16S and 23S subunits per group and sample are given. The graph reveals that the proportion of transcripts corresponding to the 23S unit is greater than that of 16S for all times except for week 24, in which the proportion of 16S transcripts is slightly greater. The ratios of 16S to 23S transcripts is shown in Figure 4. The mean ratio increases in the first week of treatment, showing a decrease in week 4, with its average being close to that estimated for the point of diagnosis. A pronounced increase is observed in week 24. The results of the Wilcoxon tests comparing the 16S/23S ratio at diagnosis with the subsequent timepoints (Table 1) indicate that the parameter is significantly greater in week 24 than prior to treatment, with mean 0.97 at diagnosis vs. 1.17 in week 24 (adj. p-value 0.027). Moreover, no significant differences in the ratio were found between diagnosis and week 1 or week 4 of the treatment.

**Table 1.** Comparisons of 16S/23S ratio at diagnosis with subsequent timepoints.

| Comparison | Statistic | p-value | Alternative | FDR p-value |
| --- | --- | --- | --- | --- |
| Diagnosis - Week 1 | 1445.0 | 0.254 | less | 0.255 |
| Diagnosis - Week 4 | 1405.0 | 0.196 | less | 0.255 |
| Diagnosis - Week 24 | 1097.0 | 0.009 | less | 0.027* |

**Figure 4.**
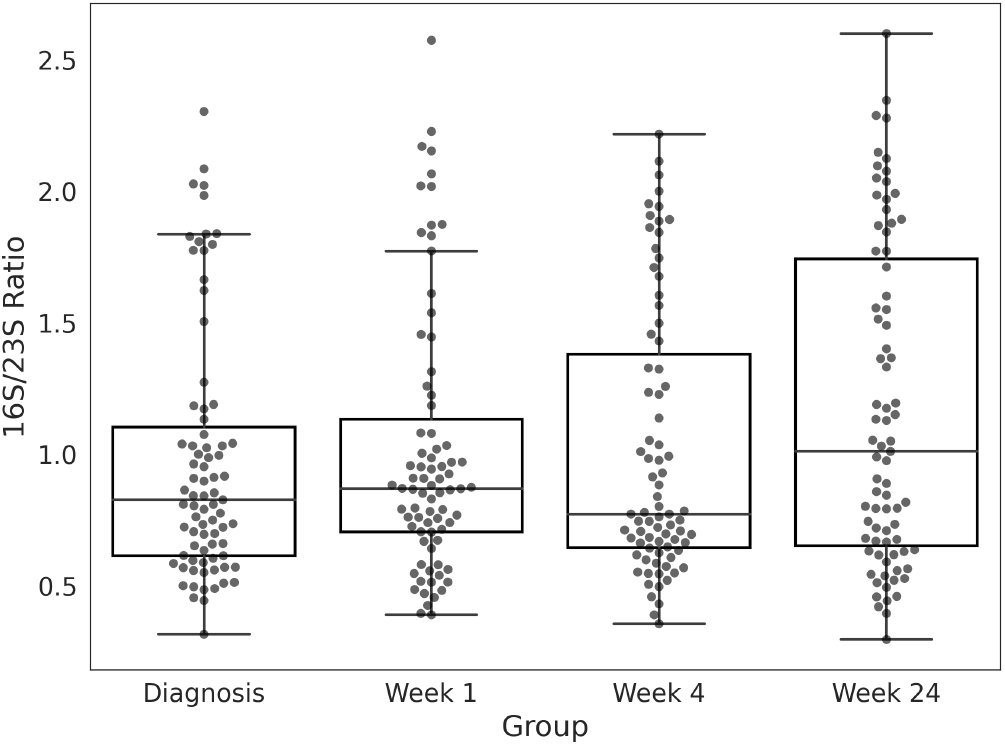
Bar-swarm plot” of 16S/23S ratios per treatment time.

Regarding the smaller units of the rRNA operon, as mentioned, they were not detected in all samples and ITS2 was not detected for any sample. Figure 3.b depicts the scaled proportions of transcripts found for 5S and ITS1 in each group. The mean relative proportion of these transcripts was higher in the time of diagnosis compared to all subsequent time points, showing a trend towards reduction during treatment. Nevertheless, as patent on the figure, a considerable inter-sample variability was observed within each group, being maximal at diagnosis.

Regarding the smaller units of the rRNA operon, as mentioned, they were not detected in all samples and ITS2 was not detected for any sample. Figure N depicts the scaled proportions of transcripts found for 5S and ITS1 in each group. The mean relative proportion of these transcripts was higher in the time of diagnosis compared to all subsequent time points, showing a trend towards reduction during treatment. Nevertheless, a considerable inter-sample variability was observed within each group, being maximal at diagnosis.

#### 0.1 Alpha and beta diversities in the course of the treatment

The Shannon index of the taxonomic distribution obtained with Bracken was estimated at the four time points (Fig. 5). A trend toward increasing the median alpha diversity in the course of the treatment is apparent on the graph.

**Figure 5.**
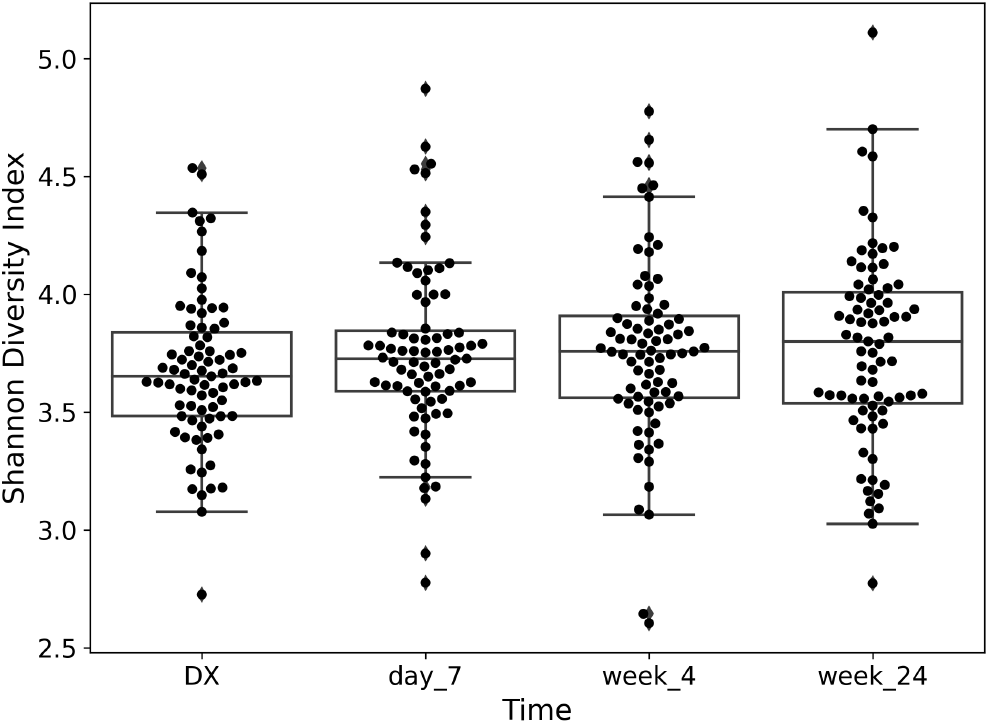
Distribution of computed Shannon indices per treatment time.

**Figure 6.**
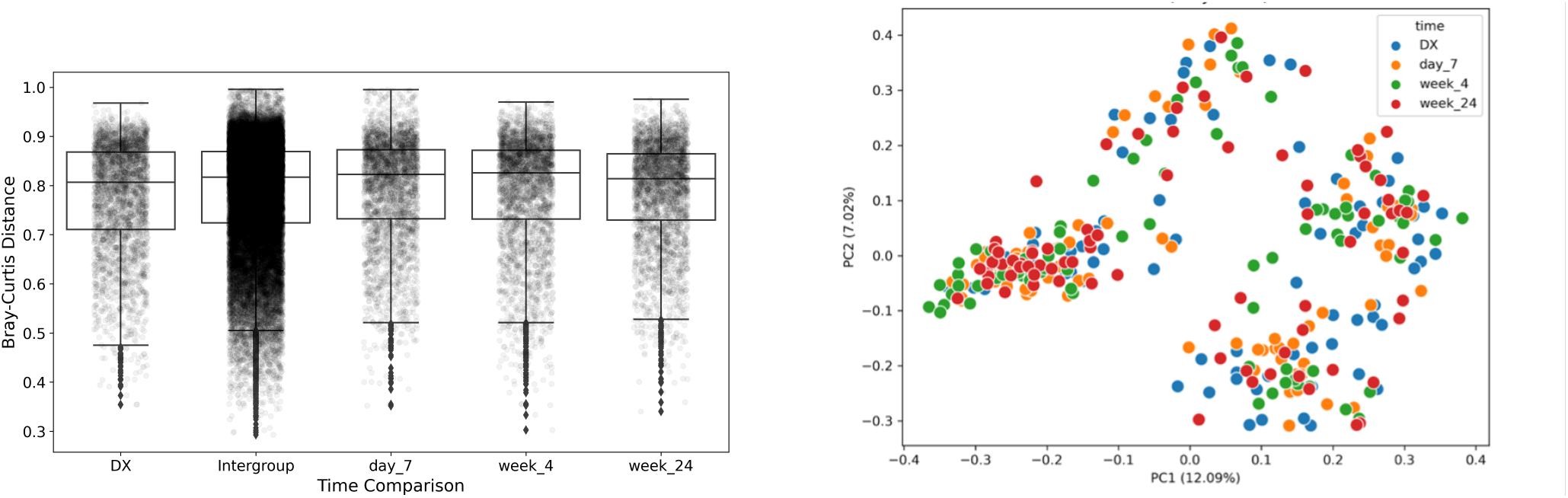
a) Proportion of 16S and 23S transcripts per treatment time and b) Scaled proportions of 5S and ITS1 subunitsfor each timepoint

The comparisons between diversity at diagnosis and subsequent sampling times applying the Wilcoxon signed-rank test are given in Table 2. Alpha diversity seems to increase in all cases, although the differences are only significant for week 1 and week 4.

**Table 2.** Alpha diversity comparison between diagnosis and subsequent treatment times.

| Comparison | Statistic | p-value | Alternative | FDR p-value |
| --- | --- | --- | --- | --- |
| Diagnosis - Week 1 | 1203 | 0.032 | less | 0.049* |
| Diagnosis - Week 4 | 1176 | 0.024 | less | 0.049* |
| Diagnosis - Week 24 | 1253 | 0.055 | less | 0.055 |

In the case of beta diversity quantified through the Bray-Curtis distances (Fig. 5.a), the application of PERMANOVA stratified by patient yielded no significant differences between times, revealing no significant differences in beta diversity across time points (F = 0.844, R^2^ = 0.008, p = 0.501). In addition, the variance explained when visualizing the average beta distances with Principal Coordinates Analysis (PCoA) in Fig 5.b is rather low (12.09% + 7.02% = 19.11%) failing the principal axes to capture differences between times.

The Wilcoxon comparison (table 3) of median beta diversity reveals very significant differences in the microbiological composition between diagnosis and follow-up times.

**Table 3.** Bray-Curtis median distances comparisons between diagnosis and subsequent treatment times.

| Comparison | Median distance | Statistic | p-value | Alternative | FDR p-value |
| --- | --- | --- | --- | --- | --- |
| Diagnosis - Week 1 | 0.744 | 3160 | 5.7e-15 | less | 5.7e-15*** |
| Diagnosis - Week 4 | 0.741 | 3160 | 5.7e-15 | less | 5.7e-15*** |
| Diagnosis - Week 24 | 0.795 | 3160 | 5.7e-15 | less | 5.7e-15*** |

**Table 4.** q-value of Mtb differential abundance across treatment times.

|  | Diagnosis vs Week 1 | Diagnosis vs Week 4 | Diagnosis vs Week 24 |
| --- | --- | --- | --- |
| q-value | 0.999 | 0.963 | 0.948 |

### Differential abundance analysis

The application of ANCOM-BC2 yielded no significant differences in MTBC abundance between diagnosis and the treatmen timepoints (table X)

## Discussion

In this study, we explore the potential of pathogen-derived rRNA transcriptomic profiles in peripheral blood as dynamic biomarkers for monitoring *Mycobacterium tuberculosis* (Mtb) response to standard anti-TB therapy. Our findings reveal several key insights into the evolution of Mtb rRNA expression signatures and taxonomic diversity throughout the course of treatment.

### Distinct patterns of rRNA transcripts distribution

Despite the technical limitations imposed by the use of host-repurposed, poly(A) enriched RNA-seq data, we found a consistent detection of 16S and 23S rRNA transcripts across all timepoints. This seems to confirm the persistence of Mtb genetic material in blood samples prior and after treatment, which confirms the findings of Gu et al., 2025. Interestingly, the relative abundance of the 23S transcript exceeded that of 16S at all timepoints except week 24, in which a modest predominance of 16S was observed.

The shift in the 16S/23S ratio, although not statistically significant after correcting for false discovery rate, seemed to increase at week 1, return to baseline by week 4, and rise sharply by week 24. This last effect was found to be significant, what suggests that these rRNA units may serve as sensitive indicators of changes in Mtb metabolic state or ribosomal activity in response to therapy. Again, this later assertion should be taken cautiously due to the nature of our data.

At diagnosis and early stages of treatment, the proportion of reads corresponding to the 23S was higher than those of 16S. Under favorable growth conditions, balanced synthesis of both subunits ensures efficient translation, and the differences observed may reflect the higher chances of 23S partial reads to be detected owing to its relatively longer span over 16S. During nutrient starvation, oxidative stress, or antibiotic exposure, ribosome synthesis can be downregulated or disrupted, being the 23S subunit more prone to degradation than 16S (Ignatov et a. 2015) The significant shift in the 16S/23S ratio by week 24 compared to earlier stages may indicate a physiological transition of the bacillary population toward a non-replicative or dormant state. In particular, in our study we observe the same trend in the shifting ratio between 16S and 23S subunits as Salina et al. (2019) when studying the resuscitation of dormant *non-culturable* Mtb although in reverse direction. In our case we may be observing the transition from a viable population to the remnants of a non-viable or dormant one at end of therapy. A central concern in interpreting rRNA-based biomarkers is the extent to which rRNA reflects bacterial viability. rRNA is more stable than mRNA and may persist beyond cell death, especially in complex biological matrices like blood (Keer & Birch, 2003). However, several studies have shown that rRNA decay correlates with bacterial viability in controlled settings (Bown et al., 2021), and that specific rRNA features—such as precursor-to-mature ratios—can distinguish between active and inactive bacterial populations (Walter et al., 2021; Honeyborne et al., 2014).

In our case, the reproducibility and structured dynamics of rRNA ratio changes argue against the random presence of degraded RNA and suggest a biological signal. Still, without direct measures of RNA integrity (e.g., through RT-qPCR targeting precursor rRNA or use of viability stains), we cannot conclusively determine whether the transcripts arise from live bacteria. Furthermore, the absence of significant shifts in *M. tuberculosis* abundance via metagenomic differential analysis (ANCOM-BC2) may reflect either biological stability or the limits of detection due to low pathogen read counts (Gu et al., 2023).

### Smaller rRNA subunits and inter-sample variability

Although showing high variability, the sporadic detection of 5S and ITS1 transcripts, more frequent in diagnosis and early stages of treatment, coupled with the complete absence of ITS2, indicates either low expression, rapid degradation, or technical limitations in capturing the shorter rRNA units from whole blood RNA-seq. The decreasing trend in the proportion of 5S and ITS1 transcripts during treatment, with maximum variability at diagnosis, could suggest more stable bacillary population, although no statistical significance can support this claim with our data. High inter-sample variability highlights the complexity of blood-based pathogen transcriptomics and suggests the need for optimized protocols or targeted enrichment strategies to improve detection of low-abundance transcripts.

### Alpha and beta diversity observations

The modest but increase in alpha diversity (Shannon index) during treatment might imply shifts in the overall taxonomic composition of the blood microbiome. In contrast, the PERMANOVA test revealed no significant differences in the compositional structure of the microbial communities between groups when estimated from a global perspective. Wilcoxon tests, however, revealed significant changes in microbiological diversity (Bray-Curtis distances) between baseline and subsequent timepoints,. The pairwise differences in beta diversity indicate that microbial community composition evolves over time, potentially driven by treatment effects or immune modulation. Such changes, however, did not translate to the visualization with PCoA, which explained a rather limited variance in its first two axes, what underscores the high dimensionality and complexity of the underlying taxonomic data, a challenge also noted in previous studies of host-pathogen transcriptomics.

### Limitations

Our study is subject to several limitations, including modest sample size (n=79) and reliance on datasets obtained from the host and subject of rRNA depletion, what limits sequencing depth and variability in non-host data. In addition, the few number of healthy controls in the study considered imepede the internal reference of our findings. In this line, this study suffers also from the lack of external validation datasets.

### Lack of differential abundance in MTBC by ANCOMBC2

In spite of the changes observed in rRNA transcript distributions and diversity metrics, differential abundance analysis using ANCOMBC2 did not reveal significant changes in overall MTBC abundance across timepoints. This could be due to high within-group variability, limited sensitivity of metagenomic approaches for quantifying low-abundance pathogens in blood, or persistent detection of Mtb nucleic acids despite declining viability. Alternatively, it may be a reflection of the limitations of the data considered characterized for a low number of non-host reads.

### Implications and future directions

Collectively, our results demonstrate the feasibility of detecting and quantifying Mtb rRNA transcripts in blood and reveal dynamic changes in transcript profiles during therapy. While total Mtb abundance did not significantly change by differential abundance testing, the evolution of 16S/23S ratio and changes in minor rRNA units provide a promising signal for future biomarker development. These findings support the hypothesis that pathogen transcriptomic signatures from peripheral biofluids, like blood, could complement current tools for monitoring treatment response, potentially enabling earlier detection of treatment failure or relapse. In this line, it clearly supports the potential adaptation of pathogen-based molecular biomarkers such as the RS ratio and the MBLA, to evaluate treatment progress from peripheral fluids. However, further work is required to validate these potential biomarkers in larger and more diverse cohorts, clarify their biological underpinnings, and assess their performance relative to established microbiological and clinical endpoints. A critical/key factor to explore is the proper sample handling and processing, focused in the determination of pathogen related transcriptomic signatures.

## Conclusions

Our study demonstrates the feasibility of detecting and quantifying Mycobacterium tuberculosis Complex rRNA transcripts in blood samples from TB patients at different treatment stages through the transcriptomic analysis of non-host data. We observed specific patterns in the relative abundance of Mtb 16S and 23S transcripts. At early stages the reads from 23S predominated over 16S, whereas at week 24 the ratio changed significantly becoming 16S transcripts more abundant than 23S. This could reflect bacterial adaptations or distinct physiological states under treatment stress, as the patterns we found seem to replicate those described in *in vitro* studies concerned with the shifting distribution of rRNA subunits in Mtb dormant cells.

Although no significant changes in the overall abundance of the MTBC complex were detected though differential abundance analysis (ANCOM-BC2), the variations in the distribution of rRNA subunits transcripts along with the changes in diversity indicate that a pathogen-based transcriptomic biomarker based on our findings deserves a deeper future exploration, especially as it might bring complementary information to currently available diagnostic and monitoring tools.

## Supporting information

Summary of 16S to 23S ratio

